# Integrating across neuroimaging modalities boosts prediction accuracy of cognitive ability

**DOI:** 10.1101/2020.09.01.278747

**Authors:** Javier Rasero, Amy Isabella Sentis, Fang-Cheng Yeh, Timothy Verstynen

## Abstract

Variation in cognitive ability arises from subtle differences in underlying neural architectural properties. Understanding and predicting individual variability in cognition from the differences in brain networks requires harnessing the unique variance captured by different neuroimaging modalities. Here we adopted a multi-level machine learning approach that combines diffusion, functional, and structural MRI data from the Human Connectome Project (N=1050) to provide unitary prediction models of various cognitive abilities: global cognitive function, fluid intelligence, crystallized intelligence, impulsivity, spatial orientation, verbal episodic memory and sustained attention. Out-of-sample predictions of each cognitive score were first generated using a sparsity-constrained principal component regression on individual neuroimaging modalities. These individual predictions were then aggregated and submitted to a LASSO estimator that removed redundant variability across channels. This stacked prediction led to a significant improvement in accuracy, relative to the best single modality predictions (approximately 1% to 4% boost in variance explained), across a majority of the cognitive abilities tested. Further analysis found that diffusion and brain surface properties contribute the most to the predictive power. Our findings establish a lower bound to predict individual differences in cognition using multiple neuroimaging measures of brain architecture, both structural and functional, quantify the relative predictive power of the different imaging modalities, and reveal how each modality provides unique and complementary information about individual differences in cognitive function.

**Author summary:** Cognition is a complex and interconnected process whose underlying mechanisms are still unclear. In order to unravel this question, studies usually look at one neuroimaging modality (e.g. functional MRI) and associate the observed brain properties with individual differences in cognitive performance. However, this approach is limiting because it fails to incorporate other sources of brain information and does not generalize well to new data. Here we tackled both problems by using out-of-sample testing and a multi-level learning approach that can efficiently integrate across simultaneous brain measurements. We tested this scenario by evaluating individual differences across several cognitive domains, using five measures that represent morphological, functional and structural aspects of the brain network architecture. We predicted individual cognitive differences using each brain property group separately and then stacked these predictions, forming a new matrix with as many columns as separate brain measurements, that was then fit using a regularized regression model that isolated unique information among modalities and substantially helped enhance prediction accuracy across most of the cognitive domains. This holistic approach provides a framework for capturing non-redundant variability across different imaging modalities, opening a window to easily incorporate more sources of brain information to further understand cognitive function.

## Introduction

Cognitive abilities are not modularly localized to individual brain areas, but rely on complex operations that are distributed across disparate brain systems (e.g., [1]). Prior work on the association between macroscopic brain systems and individual differences in cognitive ability has, by and large, relied on correlational analyses that usually assess linear changes in a particular cognitive task or measure (e.g., general intelligence quotient) that coincide with specific brain properties such as region size [2, 3], gray matter [4] and white matter [5] volume, cortical thickness [6] and surface area [7], resting-state functional connectivity [8], task-related activity [9], global functional network properties [10], white matter connectivity [11], and other unimodal measures. However, these correlation approaches, based on unimodal imaging methods, suffer several critical limitations. First, due to the mass univariate nature of the analyses, a large number of statistical tests is usually performed, thereby raising the chances of Type I error (false positives) and decreasing the statistical power of the study after adjusting for multiple testing. Second, they do not take into account the mutual dependencies between brain features and therefore ignore redundant sources of variability. Finally, the lack of out-of-sample validation tests leads to over-optimistic results (i.e., potential overfitting), thus lowering their generalizability across studies and applicability in clinical routines.

To address some of these limitations, recent studies have adopted machine learning frameworks that can accommodate all of these deficiencies by building predictive models from multivariate features across the whole brain and testing them on independent hold-out data samples. These methodologies have been widely applied to predict cognitive performance (see [12] and references therein) in out-of-sample test sets and have proven particularly popular with resting-state functional connectivity paradigms due to their inherent multivariate nature. For example, recent studies show that functional connectivity profiles, distributed across the brain, can predict up to 20% of the variance in general intelligence [13] and 25% in fluid intelligence, with regions within the frontoparietal network displaying a positive correlation and regions in the default mode network an anti-correlation [14]. Similar sparse regression models have shown how variability in white matter integrity of association pathways can reliably predict individual differences in cognitive ability [15]. By building predictive models that can be evaluated in out-of-sample test sets, as opposed to simple association analyses, these machine learning approaches can quantify the degree of generalizability of particular findings, providing key insights into potential for neuroimaging based biomarkers for cognitive function.

Despite the success of applying predictive modeling approaches for the mapping of brain properties to differences in cognitive performance, previous work has largely focused on unimodal methods. Thus, an implicit assumption is that an individual neuroimaging modality is sufficient to capture all, or at least, enough aspects of underlying neural tissue to be a reliable measure of general brain properties. Yet we know that different neuroimaging modalities reveal fundamentally different properties of underlying neural tissue. For example, functional MRI (fMRI) and diffusion MRI (dMRI) reveal separate, but complementary, properties of the underlying connectome that independently associate with different aspects of cognition [16]. This means that different measures of brain structure and function may reveal complementary associations with cognitive abilities that collectively boost the power of predictive models.

One of the challenges of building multimodal models of individual differences is the increased complexity of the explanatory model when one attempts to combine all the sources of variation. Modeling variability from a single neuroimaging modality is an already high dimensional statistical problem [17–19], with many more features than observations. Adding more modalities exponentially increases model complexity, increasing the risk of overfitting, even when traditional approaches to dimensionality reduction (e.g., principal component regression) or sparse feature selection (e.g., LASSO regression) are applied. However, one way around this dimensionality problem is transmodal learning [20], a multi-modal predictive approach that combines elements from transfer [21] and stacking (sometimes also called stacked generalization) [22] learning paradigms. Transmodal learning takes independent predictions from separate channels (e.g., generated from separate imaging modalities) and runs a second model using the single-channel predictions as the inputs. This second “stacked” model attempts to find unique sources of variability in the different input channels. Redundancy in variance, i.e., if two different imaging modalities are picking up on the same sources of variability, is accounted for through the use of feature selection methods. The end result is a more holistic prediction model that tries to explain more variance than individual input channels. Such a transmodal learning approach was recently shown to be effective at integrating structural and functional MRI measures to generate a reliable prediction of participant age whose residuals also explained individual differences in objective cognitive impairment [23].

In the present study, we used the transmodal, or stacked, learning method to quantify the extent to which the combination of data from multiple neuroimaging modalities permits increasing predictive performance in several cognitive domains, including intelligence, sustained attention, working memory, spatial orientation and impulsivity. By using a large dataset, comprised of multiple neuroimaging measures from 1050 subjects from the Human Connectome Project [24], we demonstrate that for most cognitive domains a significant enhancement in overall prediction utility is achieved when multiple modalities are integrated together, thus indicating that each brain measurement provides unique information about the underlying neural substrates relevant for cognitive function. In addition, this analysis yields a multi-modal cognitive phenotype for each cognitive ability, namely, the subset of architectural brain features whose combination yields a significant and complementary enhancement of the prediction performance.

## Results

Our primary goal was to see if predictions that integrate across neuroimaging modalities provide a boost to the prediction capability of individual differences in cognitive ability. For purposes of our analysis, the primary neural measures consisted of MRI-based assessments of (1) functional networks defined as Fisher’s z-transformed Pearson correlation coefficients between resting BOLD time series, (2) measures of cortical surface area, (3) cortical thickness attributes, (4) global and subcortical volumetric information and (5) local connectome features representing the voxel-wise pattern of water diffusion in white matter. Our multi-level, stacked modeling approach (see Fig 1 and Material and Methods section for details) uses a l1-constrained (LASSO) variant of principal component regression (PCR) to generate predictions of specific cognitive scores from single imaging modalities in a training set. These are referred to as *single-channel* models. To integrate across modalities, we stacked these single-channel predictions together and used them as inputs to a separate LASSO regression model that performs a weighted feature selection across channels. This is referred to as the *stacked* model and it produces a new set of predictions for cognitive scores of individuals by selecting and reweighting the individual channel predictions. Performance of the single-channel and stacked models are then evaluated by comparing the observed scores with the predicted scores in the out-of-sample sets. All models were fit on 70% of the data (training set) and tested on the remaining 30% (test set). A Monte Carlo cross-validation procedure [25] with 100 random stratified splits was employed to assess the generalization of these predictions.

**Fig 1.**
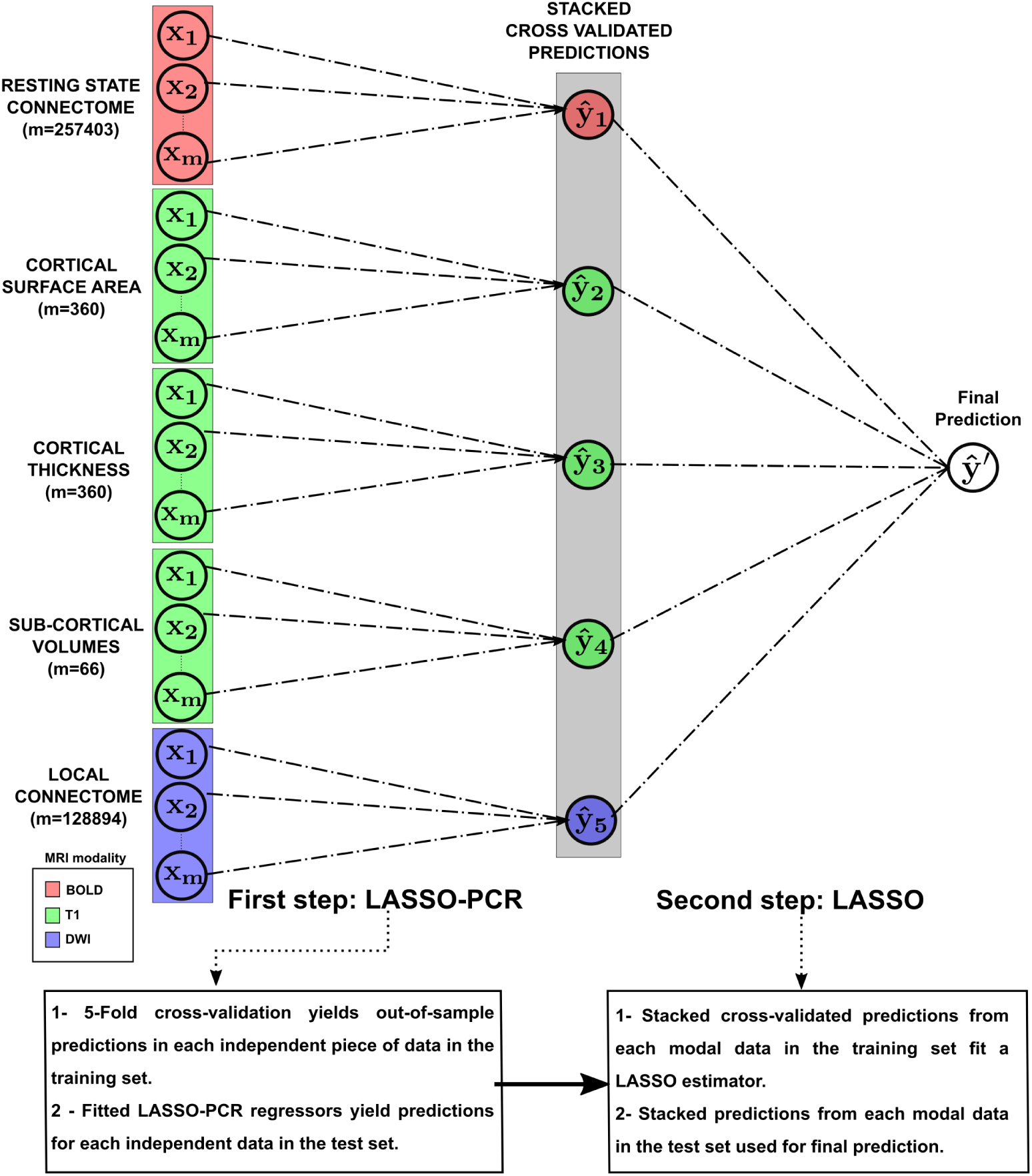
Stacking methodology for multi-modal data prediction. In the first step, a principal component regression, with an l1-sparsity (LASSO) constraint, and 5-Fold cross-validation is applied to each brain measurement to simultaneously optimize the model and generate out-of-sample predictions. These predictions are then stacked to fit a new LASSO model during the second learning step that performs weighted feature selection across single-channel predictions.

The overall performance of the single-channel and stacked models are depicted in Fig 2. These accuracies were determined using the coefficient of determination *R*^2^ (see Fig S1 for the mean absolute error scores), which shows the percent variance explained by each model in out-of-sample test sets. In Fig 3, the contributions of each channel to the stacked model (estimated by the LASSO weights) are displayed for those domains in which stacking bonus is positive, *i.e.* the scenario in which different measurements aggregate complementary and non-redundant variability. Finally, in order to understand the relative feature importance in the predictions of each brain measurement, we refit the LASSO-PCR estimator to each single-channel model using all observations. The resulting phenotype maps are depicted in Fig 4, only for those measurements whose median cross-validated contribution to the stacked model is different from zero at *α* = 0.05 significance level. In the following sub-sections we shall elaborate on the specific pattern of results for each cognitive factor represented by the scores given in Table 1.

**Table 1.** Main descriptive statistics for the response cognitive variables.

| Name of the test:<br>measure score | Mean | Median | Skewness | Kurtosis | Mild (Extreme)<br>% outliers | Lower-Higher<br>95% mean CI |
| --- | --- | --- | --- | --- | --- | --- |
| NIH Toolbox Cognition:<br>Total Composite Score | 122.28 | 120.77 | 0.21 | -0.49 | 0.00 (0.00) | 121.37-123.17 |
| NIH Toolbox Cognition:<br>Fluid Composite score | 115.43 | 114.15 | 0.26 | -0.46 | 0.00 (0.00) | 114.72 - 116.13 |
| NIH toolbox Cognition:<br>Crystallized Composite score | 117.90 | 117.81 | 0.11 | 0.09 | 0.01 (0.00) | 117.29 - 118.50 |
| Delay Discounting Test:<br>AUC for discounting<br>of \$200 | 0.26 | 0.20 | 1.33 | 1.57 | 0.04 (0.00) | 0.25 - 0.27 |
| Variable Short Penn<br>Line Orientation Test:<br>total number of<br>correct responses | 14.93 | 15.00 | -0.27 | -0.14 | 0.00 (0.00) | 14.67 - 15.23 |
| Penn Word Memory Test:<br>total number of<br>correct responses | 35.64 | 36.00 | -0.82 | 0.57 | 0.01 (0.00) | 35.45 - 35.81 |
| Short Penn Continuous<br>Performance Test:<br>sensitivity | 0.96 | 0.97 | -3.29 | 19.76 | 0.04 (0.01) | 0.95 - 0.96 |

**Fig 2.**
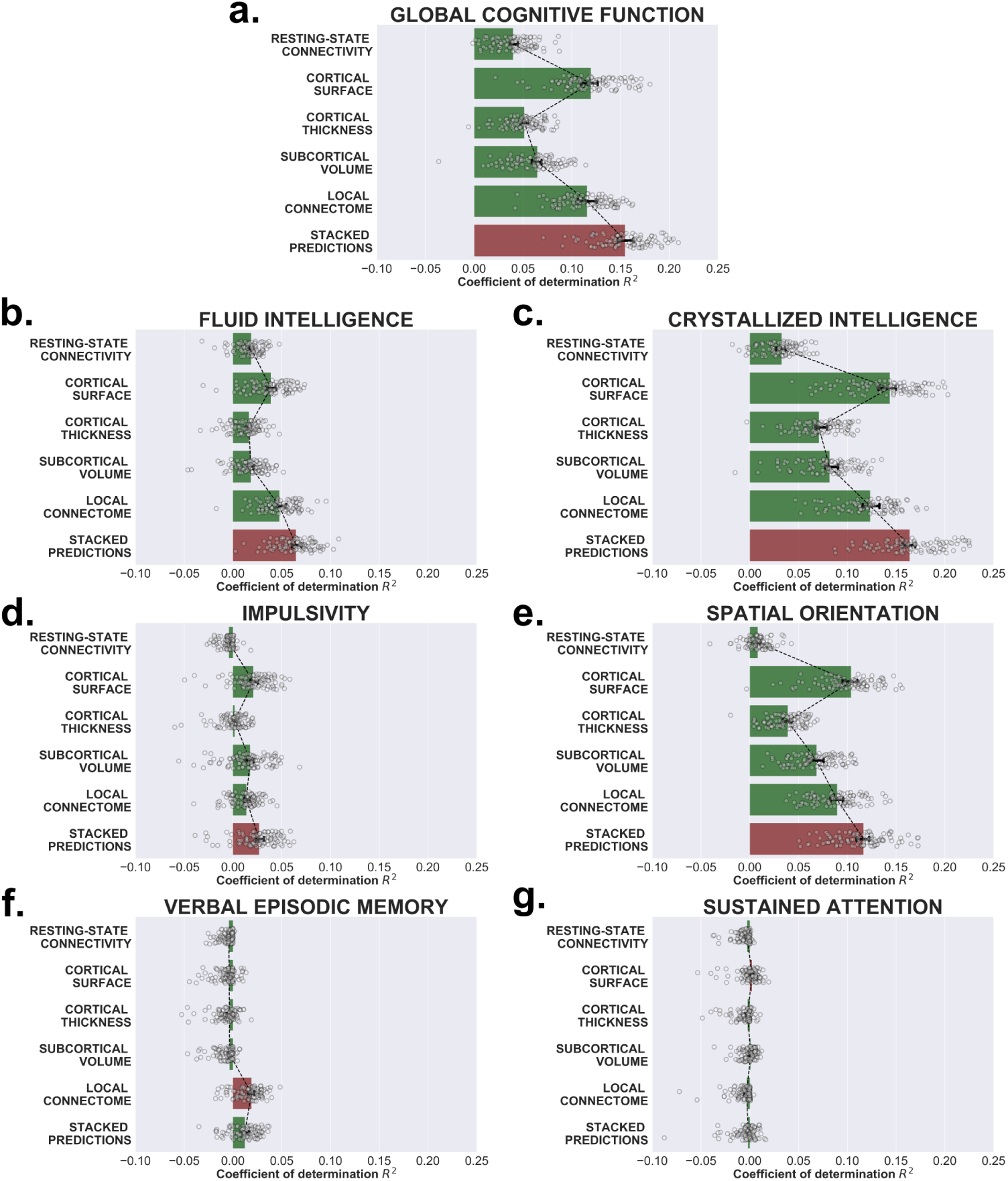
Single-channel and stacked performances to predict cognition. The coefficient of determination, *R*^2^, between the observed and predicted values of seven cognitive scores using each brain measurement separately and together by stacking their predictions. The scenario that yields the maximum predictive accuracy in out-of-sample tests is shown in red.

**Fig 3.**
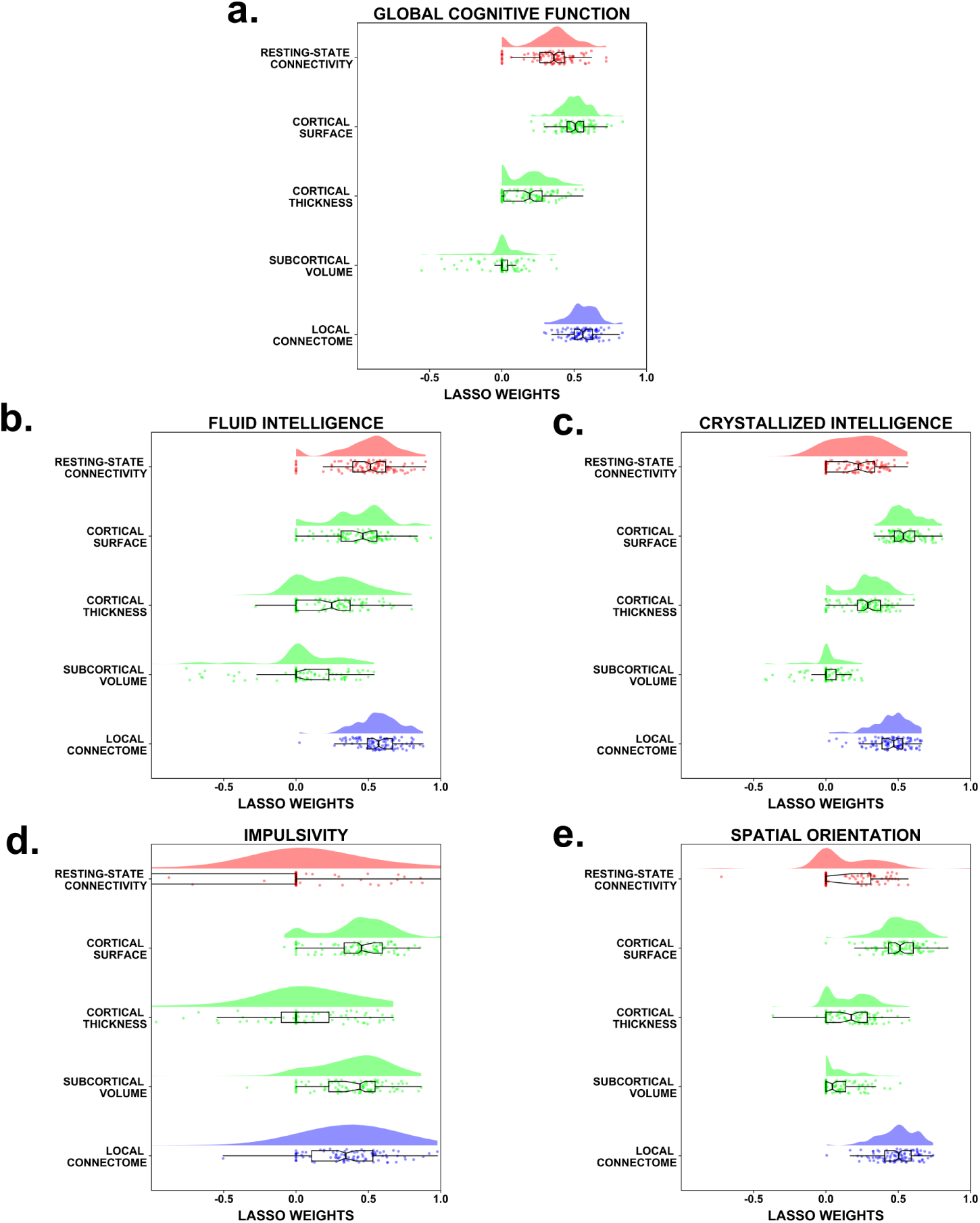
*β* coefficient distribution of each single-channel in the stacked model. Across the 100 different data splits, the weight distribution assigned to the out-of-sample predictions of each brain measurement by the stacked LASSO model in those cognitive scores in which stacking significantly improved the overall performance.

**Fig 4.**
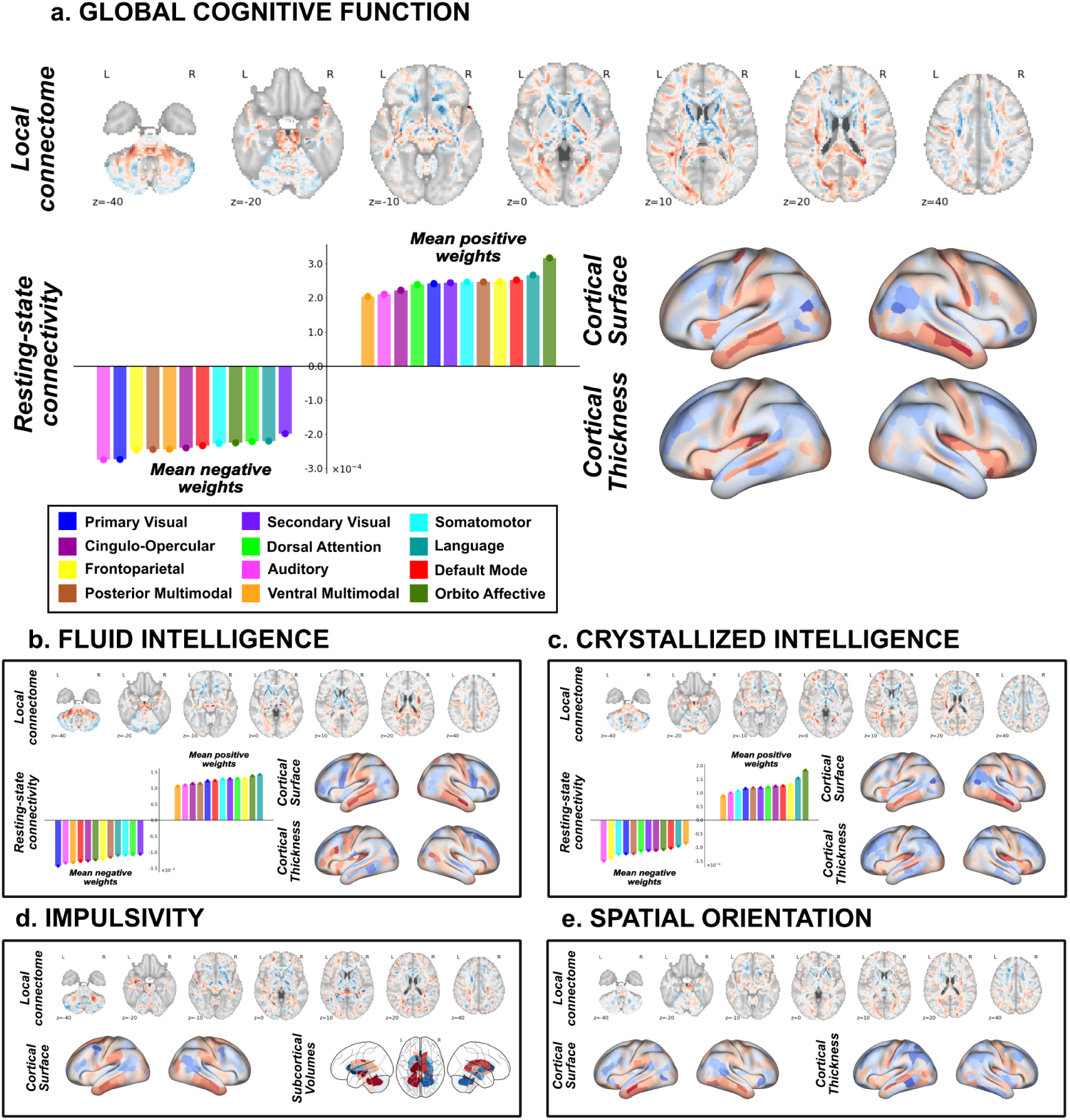
Neurophenotypes of cognitive prediction. Correlates of each brain measurement whose contribution to predicting cognitive scores during stacking is statistically non-redundant. Red and blue colors in the brain images (i.e., local connectome, cortical surface area, cortical thickness, and subcortical volumes) display positive and negative weights respectively. To facilitate interpretability, a barplot shows the average loadings of resting-state connectivity features to each intrinsic functional network (indicated by color).

### Global cognitive function

Global cognitive function was estimated by the Composite Cognitive Function score, a proxy for a general estimate of intelligence. Here the single-channel models based on cortical surface area and local connectome features produced the highest predictive rates for individual modalities, with a median *R*^2^ = 0.119, 95% CI [0.110, 0.126] and 0.116, 95% CI [0.107, 0.125] respectively. Moreover, the relative prediction accuracy of these two models did not differ statistically (one tailed Wilcoxon test *p* = 0.116, rank-biserial correlation 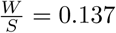). Compared to the cortical surface area and local connectome models, a significant drop in prediction performance occurred for resting-state connectivity (median *R*^2^ = 0.040, 95% CI [0.037, 0.045]), cortical thickness (median *R*^2^ = 0.051, 95% CI [0.047, 0.055]) and global and sub-cortical volumetric (median *R*^2^ = 0.065, 95% CI [0.061, 0.069]) features. Thus we see substantial variability across individual neuroimaging modalities in their predictive utility on a measure of global cognitive function.

After integrating predictions across modalities, however, an important improvement in overall accuracy is observed. The stacked model raised the median *R*^2^ to approximately 0.155, 95% CI [0.147, 0.163] in global cognitive function. Thus, the stacked model predicted a significant incremental median bonus *ℬ* = 0.035, 95% CI [0.032, 0.039] compared to the best single-channel model.

Based on the LASSO weights in the stacked model (see Fig 3a), we identified the local connectome and cortical surface areas as the strongest contributing measurements, with the former (median *β* = 0.560, 95% CI [0.534, 0.585]) contributing significantly more than the latter (median *β* = 0.508, 95% CI [0.480, 0.526]). Interestingly, resting-state connectivity was still a reliable predictor (median *β* = 0.360, 95% CI [0.337, 0.386]), as was cortical thickness (median *β* = 0.193, 95% CI [0.154, 0.223]), although to a lesser degree. A null median weight assigned to the volume channel predictions showed that these factors did not appear to reliably contribute to the stacked model. Such diverse patterns of contributions from single-channels to the stacked model may be partially affected by the L1 regularization term dealing with the shared variance between predictions. Indeed, out-of-sample predictions from volumetric factors exhibited a large collinearity, measured through Pearson correlations, especially with those predictions from cortical surface area (median *r* = 0.597, 95% CI [0.577, 0.611]) and local connectome attributes (median Pearson *r* = 0.510, 95% CI [0.499, 0.524]). In contrast, a lower similarity was found when compared to predictions from resting-state connectivity profiles (median *r <* 0.25 with respect to all measurements), hence increasing their likelihood to be present in the final predictive model.

For the largest contributing channel, the local connectome, global cognitive ability positively associated (warmer colors in Fig 4a, top panel) with signal in primarily association (i.e., intrahemispheric cortical-cortical pathways) and commissural (i.e., interhemispheric pathways) fiber systems. In contrast, many projection pathways, with the major exception being cerebellar peduncles, were negatively associated with global cognitive ability (cooler colors). This was particularly strong in the internal and external capsules. For cortical surface area, ventral temporal and anterior parietal regions areas were largely positively associated with global cognition while frontal and posterior parietal pathways were negatively associated. Interestingly, a different pattern was observed for cortical thickness, where anterior parietal regions and the insula particularly appeared to positively associate with global cognitive function. Finally, resting-state connectivity for the orbito-affective, language and default mode appeared to be the strongest contributors to positive associations with global cognition. In contrast, the auditory and primary visual networks particularly contributed negatively to global cognitive ability scores. Loadings for the volumetric features are not shown because this channel’s predictions did not survive the feature selection step in the stacked LASSO model.

### Fluid intelligence

Global cognitive ability is usually decomposed into two domains [26]: fluid intelligence (i.e., the ability to flexibly reason on new information) and crystallized intelligence (i.e., the ability to recall and use prior information). Thus we next wanted to determine how similar or different the prediction models were for these two subcomponents of general cognitive ability are. For fluid intelligence, extracted from the NIH toolbox Cognitive Fluid Composite Score, local connectome fingerprints yielded the highest coefficients of determination (median *R*^2^ = 0.048, 95% CI [0.045, 0.055]), significantly exceeding those from cortical surface areas (median *R*^2^ = 0.039, 95% CI [0.034, 0.044]), resting-state connectivity (median *R*^2^ = 0.019, 95% CI [0.016, 0.021]), subcortical and global volume features (median *R*^2^ = 0.018, 95% CI [0.015, 0.022]) and thickness of cortical regions (median *R*^2^ = 0.016, 95% CI [0.013, 0.018]). Stacked predictions raised the variability explained to a median *R*^2^ = 0.065, 95 % CI [0.060, 0.070], which is translated into a 0.015, 95 % CI [0.013, 0.019] of expected stacking bonus *ℬ*.

Interestingly, despite its weak correlation with cognitive performance, the resting-state connectivity channel supplied a non-redundant variability to the stacked model, occupying second place in variance importance (median *β* = 0.514, 95%CI [0.474, 0.560]), between the local connectome (*β* = 0.571, 95% CI [0.535, 0.610]) and cortical surface area factors (*β* = 0.462, 95% CI [0.410, 0.516]). Contributions from cortical thickness attributes were lower compared to the aforementioned channels (median *β* = 0.247, 95% CI [0.163, 0.297]). On the other hand, volumetric information appeared again to provide a null contribution to the stacked model (see Fig 3b).

Since fluid intelligence is one of the two components of the global cognitive ability score, it is not surprising that the weight maps in both domains showed large correlations across the five modalities (see Fig S3). Regardless of this, subtle phenotypic differences can still be observed (see Fig 4b); namely, the emergence of a stronger positive association with brain stem pathways such as rubrospinal tract (see Table S1) and the enhanced positive correlation of resting-state connectivity links to language areas (see also FigS2b). Loadings for global and subcortical volumes are not shown again because they did not survive the feature selection step in the stacked LASSO model.

### Crystallized intelligence

For crystallized intelligence we extracted the NIH toolbox Cognition Crystallized Composite Score. In this domain, the highest predictive accuracies were provided by cortical surface attributes (median *R*^2^ = 0.143, 95% CI [0.132, 0.149]), followed by the local connectome features (median *R*^2^ = 0.123, 95% CI [0.117, 0.133]), volumetric measurements (median *R*^2^ = 0.082, 95% CI [0.078, 0.090]), cortical thickness (median *R*^2^ = 0.071, 95% CI [0.067, 0.079]) and resting-state connectivity (median *R*^2^ = 0.033, 95% CI [0.027, 0.036]). By stacking these predictions, we obtained a median bonus *ℬ* = 0.029, 95% CI [0.025, 0.033], thus the stacked model reached a median *R*^2^ = 0.164, 95% CI [0.157, 0.170].

In contrast to global cognition, variability in the stacked model was foremostly driven by cortical surface area factors (median *β* = 0.535, 95% CI [0.510, 0.566]), which significantly exceeded the contribution from local connectome (median *β* = 0.469, 95% CI [0.431, 0.497]), cortical thickness (*β* = 0.290, 95% CI [0.260, 0.334]) and resting-state connectivity (median *β* = 0.224, 95% CI [0.148, 0.285]). Likewise, the stacked LASSO model shrunk away predictions from volume attributes when combined with the rest of channels (see Fig 3c).

Similarly to the fluid intelligence domain, multi-modal phenotypes of crystallized intelligence resembled those of global cognition, with some slight differences such as the increase of positive associations with nerve tracts between the claustrum and the insular cortex (the extreme capsule) and brainstem nerves such as the lateral lemniscus (see Table S1 and Table S2); and the enhancement of both positive and negative correlations with resting-state connectivity strength in regions of the fronto-parietal network (see Fig 4c and FigS2c).

### Impulsivity

One factor not reliably measured in the Composite Cognitive Function score is impulsivity or self-regulation, i.e., the ability to suppress contextually inappropriate behaviors. To assess this we extracted the area-under-the curve (AUC) for discounting of $200 from the Delayed Discounting Task. Even though the percent variance explained by the single-channel models for this impulsivity measure were relatively low (see Fig 2d), their performance improved using the stacked predictions (median *R*^2^= 0.027, 95% CI [0.023, 0.031]). Indeed, the stacked model performance exceeded that of the best single-channel, in this case cortical surface area properties (median *R*^2^ = 0.021, 95 % CI [0.018, 0.026]). Thus, by integrating across modalities one could expect a smallish (yet significant) median bonus *ℬ*= 0.006, 95 % CI [0.003, 0.009]. The next best single-channel performances were those from volumetric factors (median *R*^2^ = 0.017, 95% CI [0.012, 0.021]) and local connectome features (median *R*^2^ = 0.014, 95% CI [0.012, 0.017]). In contrast, both cortical thickness and resting-state connectivity metrics produced prediction accuracies with confidence intervals that are poorer than the mean response (median *R*^2^ = 0.002, 95% CI [-0.000, 0.004]; and median *R*^2^ = *−*0.004, 95% CI [-0.005, −0.003] respectively).

Interestingly, unlike the models for global cognition and intelligence, for predicting impulsivity the volumetric factor survived the LASSO feature selection step and emerged as an important explanatory source of variability in the final out-of-sample predictions (median *β* = 0.442, 95% CI [0.332, 0.477]). In fact, volumetric features explained a similar amount of variance as that of cortical surface areas (median *β* = 0.452, 95% CI [0.429, 0.534]) and more variance than the local connectome predictions (median *β* = 0.344, 95 % CI [0.294, 0.419]). On the other hand, both resting-state connectivity and cortical thickness attributes did not appear to play a role in combination with the rest of measurements (see Fig 3d).

Impulsivity phenotype maps for these contributing channels are depicted in Fig 4d. Subcortical attributes showed a strong influence of cerebellum white matter volume, an association which is manifested with opposite signs in each hemisphere. Moreover, similar loadings were also observed for the volume of the hippocampus, amygdala, pallidum, nucleus accumbens, and temporal horn. The importance of the remaining volumetric features can be found in Table S3, revealing an overall positive association of global volumetric measures with impulsivity scores. With respect to cortical surface areas, the greatest positive associations were found in areas of the dorsal-prefrontal cortex and inferior and middle temporal lobe, while negative associations were concentrated in the occipital lobe and superior temporal gyrus. Finally, particularly important areas of local connectome features positively correlated with impulsivity were found along optic radiation tracts, a group of fibers which connect the lateral geniculate nucleus of the pulvinar of the thalamus and the primary visual cortex of the occipital lobe, and brainstem pathways like the lateral lemniscus tracts (see also Table S1). In contrast, the largest negative loadings were located along the superior cerebellar peduncle, the medial lemniscus and the uncinate fasciculus tracts (see also Table S2).

### Spatial orientation

Spatial orientation is the ability to reference how the body or other objects are oriented in the environment and reflects a critical cognitive domain for spatial awareness. We extracted scores from the Variable Short Penn Line Orientation Test to look at individual differences in spatial orientation ability. The predictive models for spatial orientation followed the similar tendency observed in our other models thus far, with the stacked predictions improving overall single-channel accuracies. The best performing single-channel model was for cortical surface area (median *R*^2^ = 0.104, 95% CI [0.095, 0.110]), which was greater than the local connectome (median *R*^2^ = 0.089, 95% CI [0.083, 0.095]), volumetric measure (median *R*^2^ = 0.0689, 95% CI [0.062, 0.076]), and cortical thickness (median *R*^2^ = 0.039, 95% CI [0.034, 0.043]) models. The worst performing single-channel model for predicting spatial orientation was resting-state connectivity features (median *R*^2^ = 0.008, 95% CI [0.006, 0.011]).

Like the previous cognitive measures, the prediction of individual differences in spatial orientation improved when modalities were integrated together, with the performance of the stacked model being *R*^2^ = 0.116, 95 % CI [0.110, 0.122] and therefore constituting a stacking bonus *ℬ* = 0.016, 95 % CI [0.012, 0.019] with respect to the best single-channel model. The stacked model eliminated the contributions of resting-state connectivity and volumetric features, suggesting that these factors did not provide unique contributions to predicting spatial orientation ability above that of the cortical surface and white matter measures. As a result, only cortical surface areas (median *β* = 0.512, 95% CI [0.482, 0.545]), local connectome features (median *β* = 0.504, 95% CI [0.478, 0.527]) and cortical thickness attributes (median *β* = 0.175, 95% CI [0.100, 0.224]) appeared to provide a non-redundant contribution to the stacked model (see Fig 3e).

Finally, weight maps estimated from these contributing channels are displayed in Fig 4e. Interestingly, we can appreciate a positive correlation with local connectome features particularly along projection pathways connecting to regions in the occipital cortex (optical radiation, central tegmental and occipito-pontine tracts), which are involved in visual demanding tasks, whereas negative associations are mainly dominated by cranial nerves and axons from the rubrospinal tract. Regarding structural cortical attributes, positive loadings from surface area factors were found in regions along the inferior and middle temporal gyri and the paracentral gyrus, whereas negative associations took place in the frontal cortex. On the other hand, except for regions in the central sulci, orbital gyrus and temporal pole, thickness properties exhibited a negative correlation with spatial orientation that spans over the entire brain cortex.

### Verbal episodic memory

The Penn Word Memory test captures verbal memory abilities. For this response variable, stacked prediction accuracy (median *R*^2^ = 0.012, 95% CI [0.010, 0.016]) did not improve the single-channel’s best performance represented by the local connectome fingerprints (median *R*^2^ = 0.019, 95% CI [0.016, 0.022]). The rest of the measurements all exhibited a negative median *R*^2^, meaning that they perform worse than using predictions from the mean response variable (see Fig 2f). As a consequence, neither the *β* coefficients showing the single-channel contributions to the stacked model nor the weight maps were reported for this cognitive score.

### Sustained attention

The Short Penn Continuous Performance Test is the measure of sustained attention. As shown in Fig 2g, accuracies of single-channels and stacked model in this domain were overall negligible and worse or not statistically different from those provided by the mean response variable. Owing to this, neither the *β* coefficients showing the single-channel contributions to the stacked model nor the weight maps were reported for this cognitive score.

## Discussion

Here we tested whether multiple functional, diffusion, and morphological MRI-based measurements of brain architecture constitute complementary sources of information for predicting individual differences in cognitive ability. We accomplished this by means of a stacking approach for multimodal data, a two-level learning framework in which groups of features are first separately trained and their predicted response values subsequently stacked to learn a new model that takes into account redundant effects across channels. Our results show that for most of the cognitive measures tested integrating across different brain measurements provides a boost to prediction accuracy, highlighting how different imaging modalities provide unique information relevant to predicting differences in human cognitive ability.

One of the strengths of our approach is the assessment of how performance in different cognitive domains associates with a wide range of brain measurements. Overall, our results show that effect sizes tend to be moderate (at most explaining less than 20% of the variance), which begs the question of where the remaining variability may come from. It might be possible that the metrics employed here, which largely reflect static architectural aspects of global brain systems, are missing the fundamental dynamics of neural circuits during relevant behavioral states for expressing specific cognitive functions. In this regard, specific task-evoked fMRI measurements that directly assess the neural reactivity during cognitive evaluation [27–29] could help raise the overall predictive power. For example, we recently showed that brain activation patterns during affective information processing tasks predict an important portion of individual differences of cardiovascular disease risk factors, a finding that we could not have reached had not we used the appropriate and specific task fMRI experiment [30]. Additionally, increasing both spatial resolution to better capture features of structural-functional variation [31] and temporal resolution for a more accurate decoding of the underlying brain dynamics [32] could be valuable and complementary sources of cognitive performance correlation. Thus we consider the work here a proof-of-principle for making holistic models that predict specific cognitive abilities, which could be further improved with additional, more specific, inputs.

Of particular note is that the observed effect sizes from resting-state connectivity are consistently small, which appears to be in conflict with previous results that reported a medium-large correlation (*r* = 0.5) between patterns of resting-state connectivity and fluid intelligence functioning [14]. In our case and for this particular domain, the maximum performance that we achieved across all simulations is ostensibly smaller (*R*^2^ = 0.047, *r* = 0.29). Nevertheless, such a decrease in the effect sizes was expected due to our use of a much larger sample size (*N* = 1050 versus *N* = 126), which reduces inflated results caused by sampling variability and therefore, findings are more reproducible and inferred patterns generalizable to a broader population spectrum [33]. In addition, it is important to note that our preprocessing pipeline does not regress out the mean global BOLD signal, which is supposed to strengthen the association between resting-state functional connectivity profiles and behavior [34]. This step is still controversial since it is not clear whether it supplies real or spurious information [35]. Finally, we relied on the coefficient of determination, *R*^2^, to assess the predictive power of the learned models, in contrast to using Pearson correlation coefficient, which overestimates the association between predicted and observed values and therefore it is not a recommended performance metric during regression tasks [36, 37].

The overall stacking approach to multimodal integration that we applied here closely follows work by Liem and colleagues [23], who used a stacking approach with multimodal brain imaging data to improve the performance in individual age prediction, although with two big differences in structure. First is the choice of the algorithm for the second learning stage in this transmodal approach. Albeit performances from a Random Forest algorithm, as used by Liem and colleagues, would have been proven to be less variable compared to other well known algorithms in similar scenarios [38], we decided to use a LASSO regression model because of its simplicity (it only has one hyperparameter to tune) and due to the fact that the L1 penalty term can automatically get rid of the redundant variability of the different channels. The second difference is the number of neuroimaging modalities, since we have also included diffusion data in our study. Indeed, we have demonstrated that the inclusion of local connectome features played an important role for prediction, since they alone account for a moderate rate of variability consistent across all cognitive domains. Moreover, such variability survives and indeed prevails when combined with the rest of single-channel predictions. This finding validates the role of the local connectome fingerprint as a reliable correlate of cognitive factors at the individual level [15] and suggests its complementary role in combination with other brain measurements. In particular, white matter diffusion tracts provide a putative structural basis for the macroscopic human connectome that is reflected in the correlation for age [39] and cognition [40, 41]. Furthermore, it is important to stress that local connectome fingerprints do not rely on fiber tracking algorithms, which reduces the risk of false-positive bias when mapping white matter pathways [42–44]. Our findings here with respect to predicting cognitive ability, along with the work of Liem and colleagues on age predictions [23], simply demonstrate how powerful a stacking approach can be to maximizing explainable variance from multiple imaging modalities.

Interestingly, although stacked predictions clearly increase the variability explained at a global cognitive level, this is not the case across all domains. For example, global, fluid, and crystalized intelligence, impulsivity and spatial orientation all show improvements in prediction accuracy, in contrast to attention and word memory functioning. These results might be caused by the existence of a hierarchical cognitive categorization, with high complex functions demanding the integration of multi-modal aspects of the brain compared to lower level functions [45]. Alternatively, individual differences in some cognitive areas might be mostly parametrized by one specific brain sub-system that captures all the variability. For instance, white matter structure is an important substrate of cognitive performance whose deterioration, notably in the hippocampus, is the first sign of memory decline at both early and late stages in Azheimer’s disease [46, 47]. On the other hand, it might also happen that for certain cognitive domains, the measurements considered in this study do not constitute a sizable source of variability and therefore stacking is only aggregating noise to the predictive model. Finally, associations with cognitive performance might be affected by the inherent nature of the cognitive tests, either due to an imperfect design that adds unwanted variability, or because of the properties of the sampling distribution. For example, in our dataset, scores in the Short Penn test display a heavy deviation from normality, that can affect sensitivity in regression models.

A possible limitation of our study arises from the fact that predefined test scores were employed as response variables, which in most cases are largely coarse measures of cognitive ability that may rely on redundant underlying subprocesses, leading to a degree of similarity in the brain architecture features that contribute to predicting individual differences. Regardless of this, spatial correlation analysis shows that a portion of each cognitive measure’s weight map is unique (including for the global cognitive score that is constructed from both crystallized and fluid intelligence scores) and therefore there exist brain structures whose roles are specific to the area of cognition involved (see Fig S3). Future studies might benefit from adopting more sensitive approaches to measuring specific cognitive factors (e.g., psychophysical measurements) that can carefully isolate primary cognitive abilities.

Finally, our current framework treats brain measurements separately at a first step, based on the assumption that they represent independent and non-overlapping sources of cognitive variation. As we showed, this is far from true: there exists some degree of redundancy across the imaging modalities. Future studies might attempt to find a decomposition into multidimensional representations of unique and shared variance across brain measurements. This would likely increase the number of single-channels whose individual predictions can be later exploited by our stacking approach.

Despite these limitations, our work here builds on the growing body of work attempting to integrate information from different neural sources so as to maximize explained variability of individual differences. Our approach predicts individual differences in cognition by separately fitting measurements of structural, functional and diffusion modalities and subsequently stacking predictions to enhance overall accuracy while removing redundant contributions. Even though a large portion of variance in the data remains unaccounted for, our results demonstrate that effect sizes can be easily increased by using multimodal neuroimaging data and establish a solid and reliable lower bound for cognitive prediction in different domains.

## Materials and Methods

### Participants

We used publicly available data from the S1200 release of the Human Connectome Project (HCP) database (https://ida.loni.usc.edu/login.jsp). The HCP project (Principal Investigators: Bruce Rosen, M.D., Ph.D., Martinos Center at Massachusetts General Hospital; Arthur W. Toga, Ph.D., University of Southern California, Van J. Weeden, MD, Martinos Center at Massachusetts General Hospital) is supported by the National Institute of Dental and Craniofacial Research (NIDCR), the National Institute of Mental Health (NIMH) and the National Institute of Neurological Disorders and Stroke (NINDS). HCP is the result of efforts of co-investigators from the University of Southern California, Martinos Center for Biomedical Imaging at Massachusetts General Hospital (MGH), Washington University, and the University of Minnesota. Out of the 1200 participants released, 1050 subjects had viable T1-weighted, resting-state fMRI, and diffusion MRI data. In addition, 21 subjects were discarded due to the presence of missing information in some of the response variables used in this study. The final dataset comprised of 1029 individuals (550 female, age range 22-37, mean *± σ*_*age*_ = 22.73 *±* 3.68 years).

### Predictor variables

Preprocessing steps included spatial artifact/distortion removal, surface generation, cross-modal registration and alignment to standard space and the automatic ICA-FIX denoising of functional acquisitions, among others (more details on these and other additional preprocessing steps can be found in [48]).

*Structural* predictors were composed of cortical thickness (CT) and surface area (CS) values of 360 regions in a multi-modal parcellation [49], and 66 features containing global, subcortical and other volume (VL) information, directly extracted from the *aseg.stats* freesurfer file of each subject.

*Functional* predictors were estimated from the resting-state data by first computing the averaged time series from the voxels within each of the 718 regions in a parcellation which extends the aforementioned multi-modal atlas to include 358 subcortical regions [50]. Furthermore, this parcellation identified 12 different intrinsic functional networks, which included the well-known primary visual, secondary visual, auditory, somatomotor, cingulo-opercular, default-mode, dorsal attention and frontoparietal cognitive control networks; novel networks like the posterior multimodal, ventral multimodal, and orbito-affective networks; and the identification of a language network [50]. Next, a functional connectome for each subject was built by calculating the z-transformed Pearson correlation coefficient between pairs of time series. Finally, the upper triangular elements were extracted to form the final vector of 257403 functional connectivity (FC) features per subject.

*Diffusion* predictors were represented by the local connectome fingerprint (LC), a structural metric that quantifies the degree of connectivity between adjacent voxels within a white matter fascicle defined by the density of diffusing spins [51]. The local connectome was computed by reconstructing the spin distribution functions (SDFs) in all white matter voxels in a common template space, previously derived from 842 subjects of the HCP dataset [52], using q-space diffeomorphic reconstruction [53] and sampling the quantitative anisotropy [54] at peak directions within each voxel. This produces a fingerprint vector of 128894 fibers across the entire brain for each subject. These features were obtained using DSI Studio (http://dsi-studio.labsolver.org), an open-source toolbox for connectome analysis from diffusion imaging.

### Response variables

A subset of seven cognitive test scores available in the HCP repository were used as response variables [55]. Each of these measures assesses the individual performance in cognitive domains that are different to a greater or lesser extent. In particular, we selected: (a) the Unadjusted scale NIH Toolbox Cognition Total Composite Score, which provides a measure of *global cognitive function*, (b) The Unadjusted NIH Toolbox Cognition Fluid Composite Score for *fluid intelligence*, (c) The Unadjusted NIH Toolbox Cognition Crystallized Composite Score for *crystallized intelligence*, (d) the area-under-the-curve (AUC) for Discounting of $200 in a Delay Discounting Test for *impulsivity*, (e) the total number of correct responses in a Variable Short Penn Line Orientation Test for *spatial orientation* assessment, (f) the Total Number of Correct Responses in a Penn Word Memory Test, which aims at testing *verbal episodic memory*, and (g) sensitivity in a Short Penn Continuous Performance Test for *sustained attention* performance. The main descriptive statistics for these variables can be found in Table 1.

### Prediction Models

The prediction of each cognitive score was carried out adopting a transmodal approach to stacking learning. Stacking belongs to the ensemble paradigm in machine learning and it is based on a multi-level training in which predictions from a given set of models are combined to form a new meta feature matrix [20, 22]. This new matrix can be then fed into a new model for final predictions or passed to a successive and intermediate learning level.

Our transmodal scenario consisted of two-levels of learning and differs from usual stacking approaches in that each predictive model comes from training separately our different groups of features (called channels), each corresponding to the resting-state connectome, cortical thickness attributes, cortical surface areas, global and subcortical volumetric information and the local connectome fingerprints.

The entire prediction modeling procedure is sketched in Fig 1. First, we split the data such that 70% of the subjects were used for training and the rest for testing. Since we are dealing with unadjusted response variables, before fitting any model we regressed each cognitive score in the training set onto the individual ages and used the estimated regression coefficients to remove the age effect in both training and test set. Given the narrow range of ages of our data (22-37 years), this effect is likely to very weak, so perhaps this step could have been omitted. After age adjustment, a 5-Fold cross-validation was applied to the observations in the training set using a principal component regression model with a L1 regularization term (LASSO-PCR) on each channel. This estimator constitutes a pipeline with the following sequential steps:

1. Dimensionality reduction by PCA to the input matrix of features *X* of each channel:

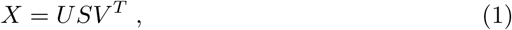

where the product matrix *Z* = *US* represents the projected values of *X* into the principal component space and *V* ^*T*^ an orthogonal matrix whose row vectors are the principal axes in feature space.
2. Regression of the (age adjusted) response variable *y* onto *Z*, where the estimation of the *β* coefficients is subject to a L1 penalty term *λ* in the objective function:

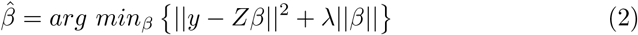
3. Projection of the fitted 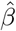 coefficients back to the original feature space to produce a weight map (or phenotype) 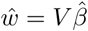 used to generate final predictions *ŷ*:

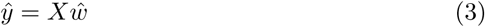

The cross-validation loop was used so as to determine the optimal value of *λ* and simultaneously generate out-of-sample predictions that are to be used at the subsequent learning level. After training each LASSO-PCR model with the optimal level of shrinkage, single-channel predictions from the test set were aggregated across all channels.

At the second level, both out-of-sample predictions from the training set and predictions on the test set were stacked across channels to form a new training set consisting of 70% observations *×* 5 channels and a new test set consisting of 30% observations × 5 channels, respectively. By means of a new *×* 5-Fold cross-validation loop, the former matrix first optimized and then fit a LASSO regression model that made predictions on the latter object of stacked data. The use of LASSO produced a feature selection as well, that automatically selected and weighted how much each channel contributed to the best final prediction.

Prediction error was measured through the coefficient of determination *R*^2^ and the mean absolute error (MAE). In order to assess the stability and consistency of the predicted metrics, a Monte Carlo cross-validation was employed. In particular, we generated 100 random data splits with the same training and test size proportions as given above (70% and 30% respectively) and repeated the entire predictive procedure for each of these splits. Thereafter, the model performance and the contributing weights of each single-channel to the LASSO stacked model were reported using the median and its bootstrapped confidence intervals at a significance level *α* = 0.05. The choice of the median was particularly suitable when summarizing the contributions of each single-channel, since the stacked LASSO model produced sparse solutions. All predictive analyses were carried out using scikit-learn [56].

### Stacking bonus

In order to formally compare the integrated model, *i.e.* the model that integrates predictions across modality, against the single-channel predictions, we defined a stacking bonus score *ℬ* which reads

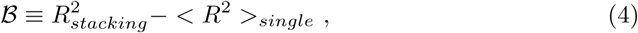

where 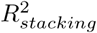 is the out-of-sample coefficient of determination from the second-level LASSO learning to the stacked predictions and 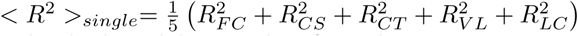 is the average performance across individual modalities. Therefore, this quantity aims to mimic the notion of synergy from information theory, which states that joint systems may convey more information that just the sum of its parts [57–59].

Since the above definition may lead to very optimistic bonuses in the presence of poor modalities, we adopted a more conservative definition by expressing this difference just against the best single-channel performance 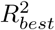 as follows

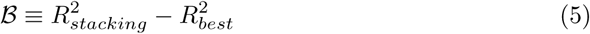

## Code and data availability

The code used to generate all the results and plots in this study is available in https://github.com/CoAxLab/multimodal-predict-cognition. The weight maps for the local connectome, cortical surface area and thickness, and sub-cortical volumetric features have been uploaded as NIFTI files to https://neurovault.org/collections/VXOBBMIZ. The data used and generated in this study are available in https://figshare.com/s/b97d2d1ba359e6458cb5.

## Acknowledgments

Data collection and sharing for this project was provided by the Human Connectome Project (HCP; Principal Investigators: Bruce Rosen, M.D., Ph.D., Arthur W. Toga, Ph.D., Van J. Weeden, MD). HCP funding was provided by the National Institute of Dental and Craniofacial Research (NIDCR), the National Institute of Mental Health (NIMH), and the National Institute of Neurological Disorders and Stroke (NINDS). HCP data are disseminated by the Laboratory of Neuro Imaging at the University of Southern California.

## Supporting information

**Fig S1.**
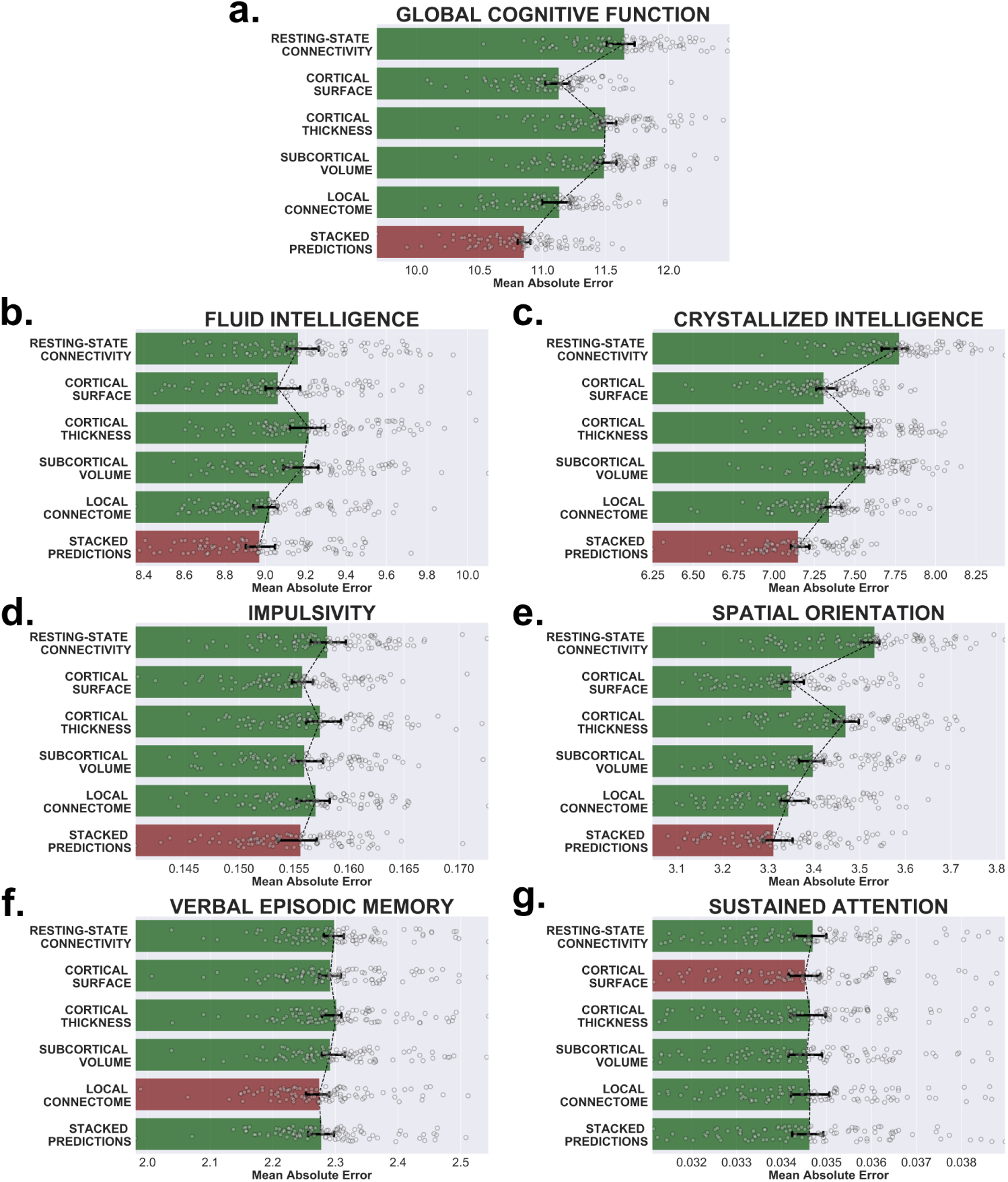
Single-channel and stacked performances to predict cognition. Mean absolute errors (MAE) between the observed and predicted values of seven cognitive scores using each brain measurements separately and together by stacking their predictions. In red the scenario that yields the maximum score.

**FigS2.**
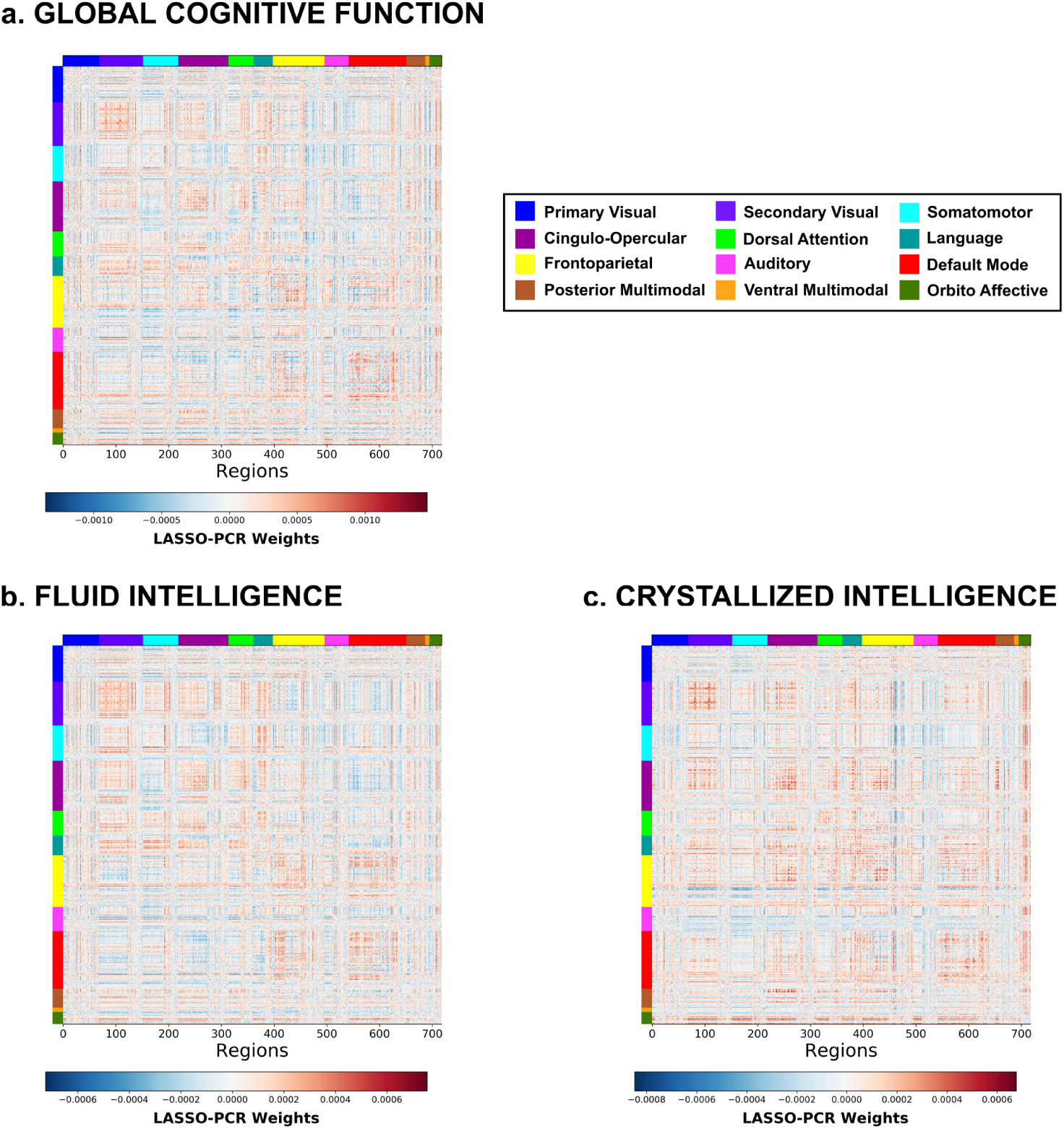
Resting-state connectivity phenotypes. Weight maps from the resting-state connectivity matrix coefficients for the three cognitive areas (global cognitive function, fluid and crystallized intelligence) in which these attributes significantly contributed to stacking. Red and blue colors display positive and negative weights respectively.

**Fig S3.**
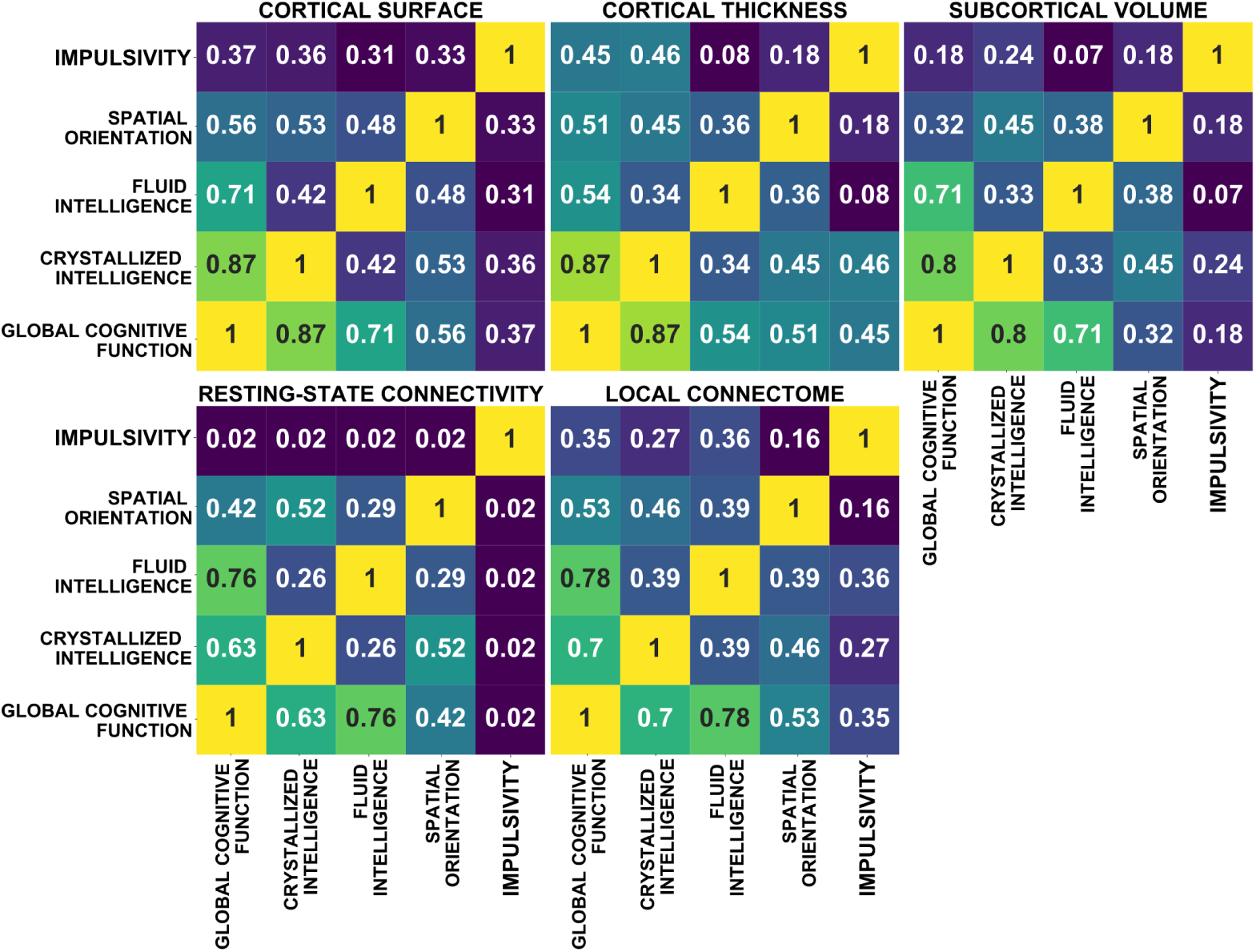
Phenotype similarity between cognitive domains. For each brain measurement, the Pearson correlation coefficient is computed to assess the similarity between the brain correlates of each cognitive score in which stacking led to a significant performance enhancement.

**Table S1.**
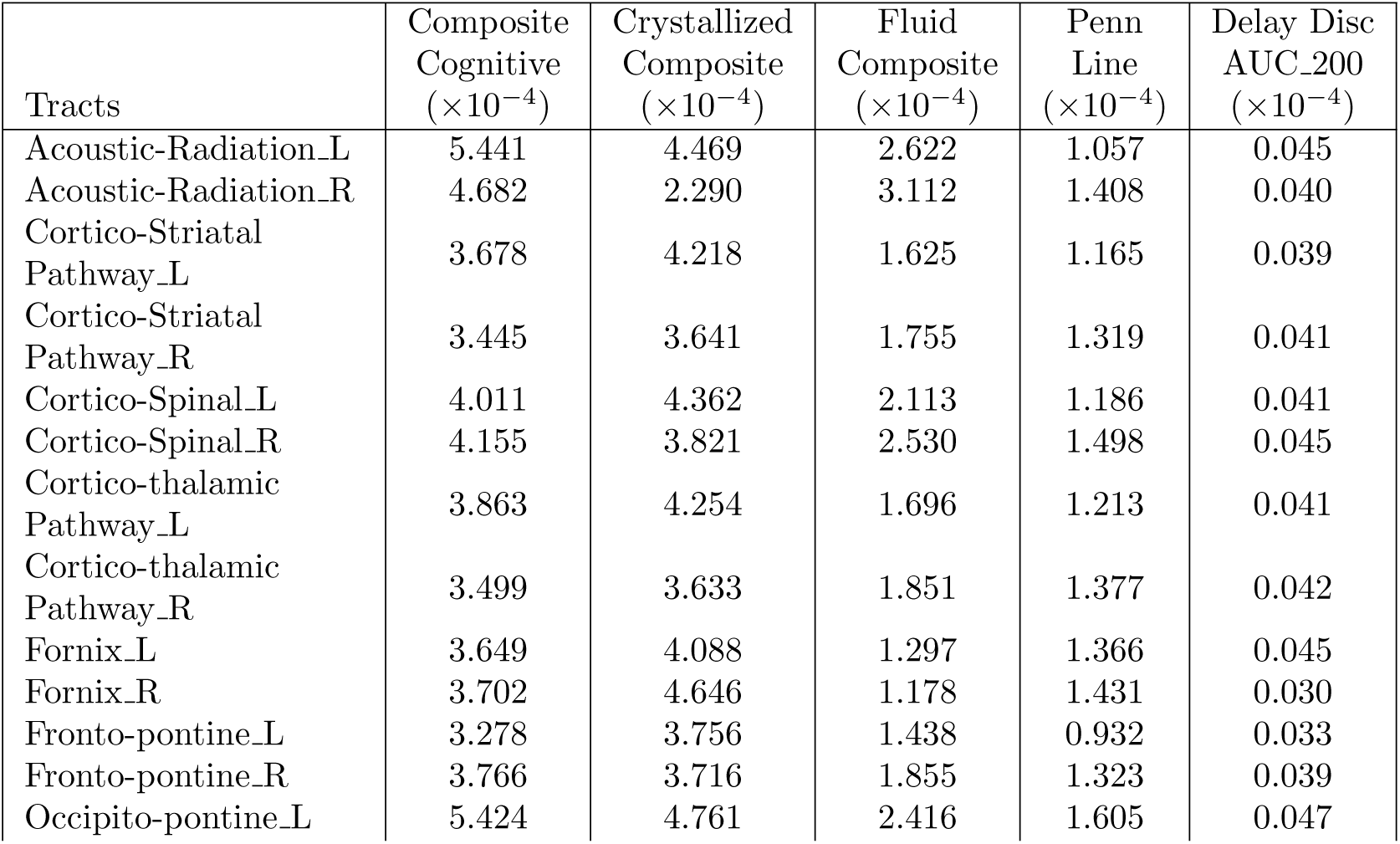
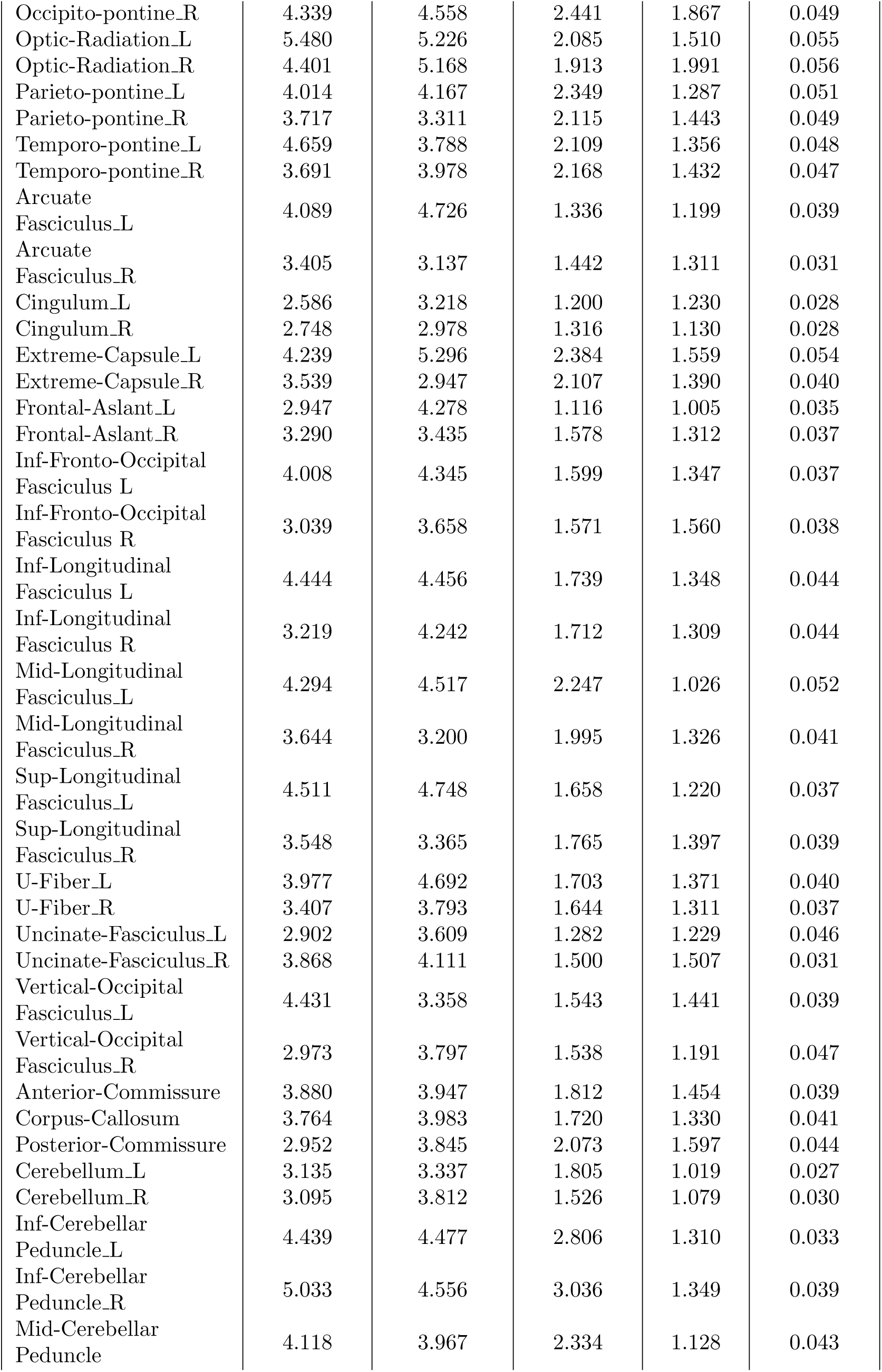
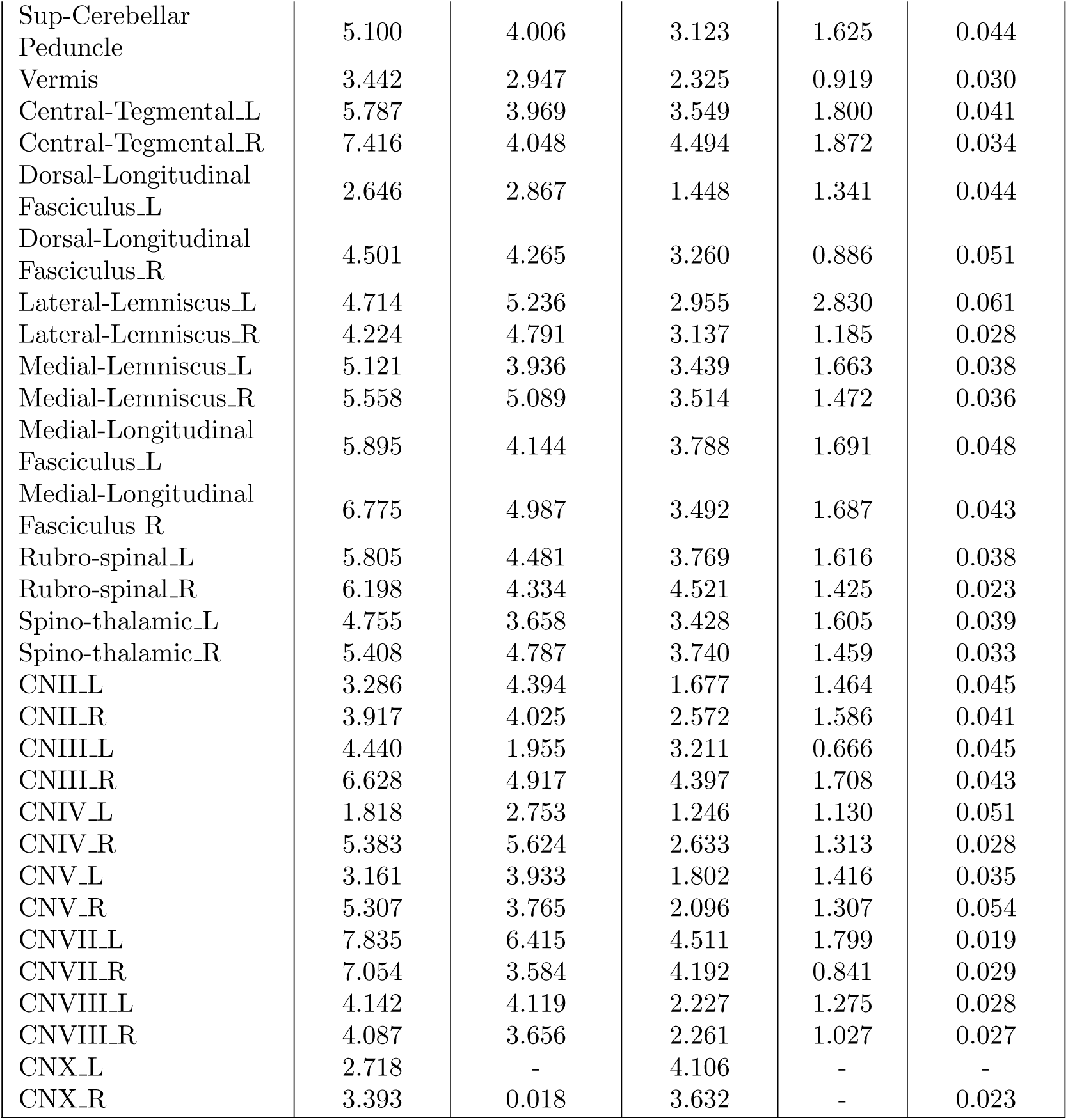
Average positive weights of local connectome features per major tract. Following a population-based atlas of structural connectome, positive loadings within each major tract have been averaged. An entry with a line mark denotes the lack of positive weights within that tract. L*≡*Left hemisphere, R*≡*Right hemisphere

**Table S2.**
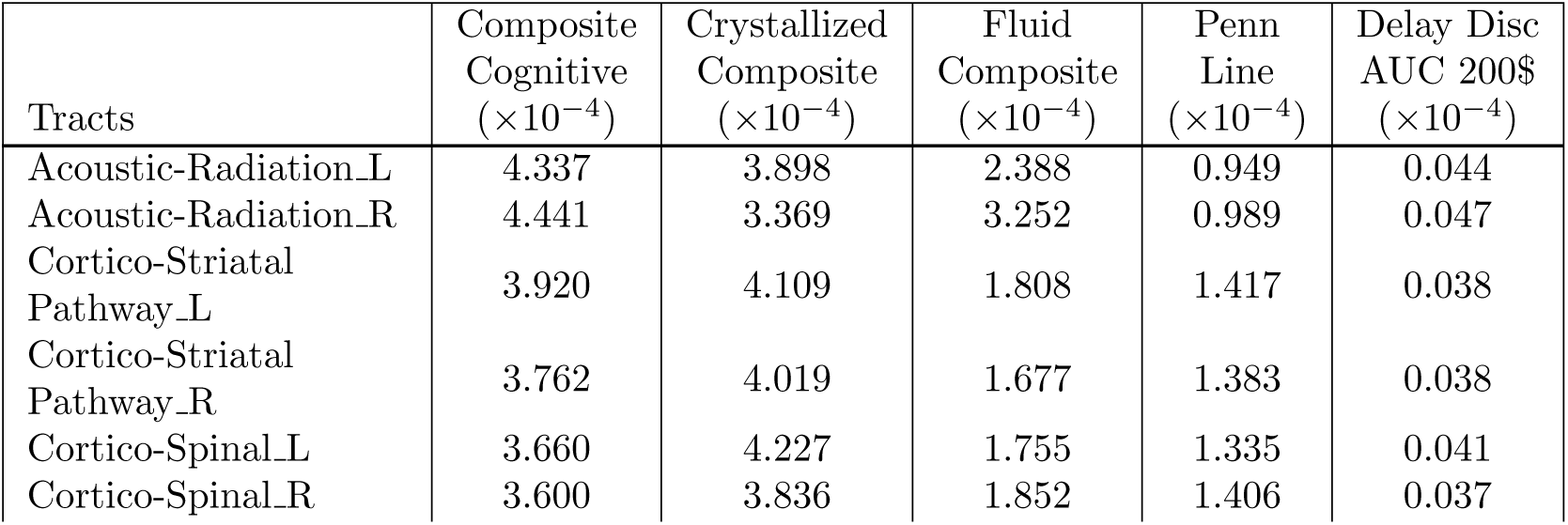
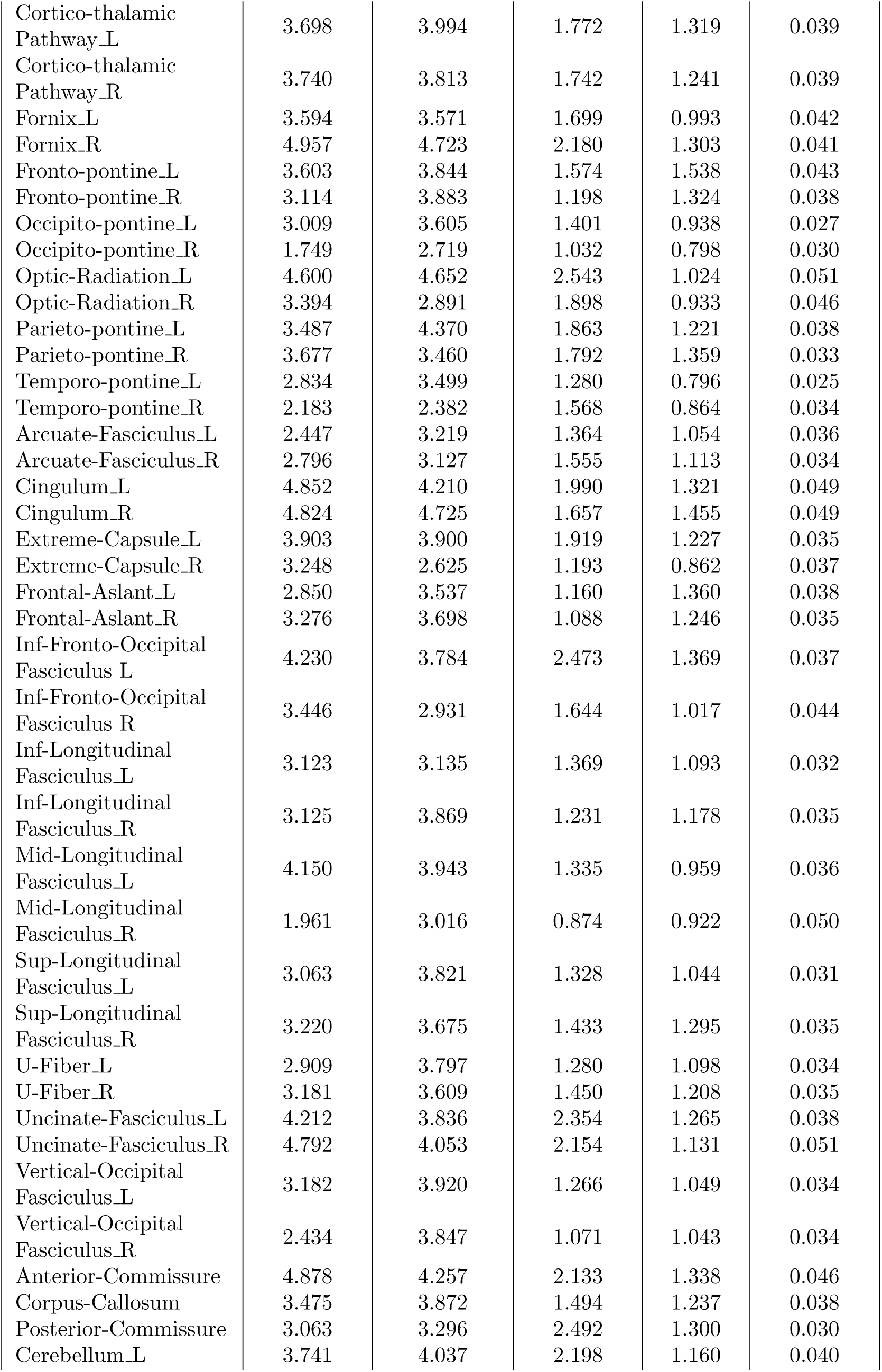
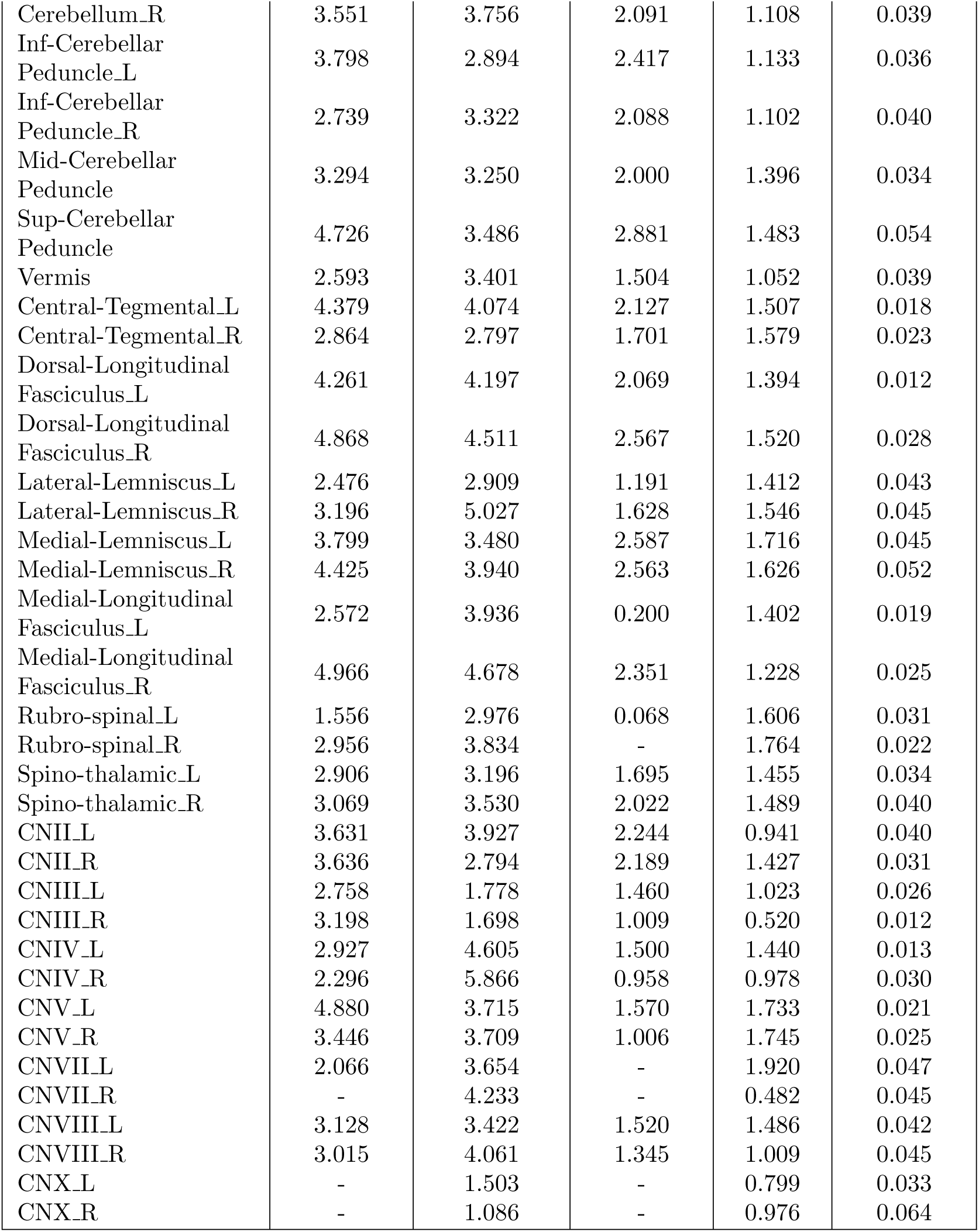
Average negative weights of local connectome features per major tract. Following a population-based atlas of structural connectome, negative loadings within each major tract have been averaged. An entry with a line mark denotes the lack of negative weights within that tract. L*≡*Left hemisphere, R*≡*Right hemisphere

**Table S3.** Weights of volumetric properties. Loadings of global and subcortical volume features for predicting the score of a Delay Discounting test, which assesses impulsivity abilities. The name of the features are the same that can be found in the *aseg.stats* file from freesurfer.

| Feature | Weight |
| --- | --- |
| rhCortexVol | 0.015890 |
| CortexVol | 0.015644 |
| lhCortexVol | 0.015265 |
| Left-VentralDC | 0.013799 |
| Left-Cerebellum-White-Matter | 0.012979 |
| MaskVol-to-eTIV | 0.012450 |
| TotalGrayVol | 0.012299 |
| CC_Mid_Posterior | 0.012187 |
| MaskVol | 0.011233 |
| Right-Lateral-Ventricle | 0.010319 |
| 4th-Ventricle | 0.008717 |
| CSF | 0.006599 |
| Right-Accumbens-area | 0.006029 |
| Right-VentralDC | 0.006001 |
| Left-Amygdala | 0.005821 |
| Left-Lateral-Ventricle | 0.004951 |
| Right-vessel | 0.004450 |
| CC_Central | 0.003990 |
| EstimatedTotalIntraCranialVol | 0.003390 |
| rhSurfaceHoles | 0.003077 |
| WM-hypointensities | 0.003074 |
| Brain-Stem | 0.002957 |
| Right-Inf-Lat-Vent | 0.002753 |
| Right-Caudate | 0.002452 |
| Left-Caudate | 0.002396 |
| CC_Mid_Anterior | 0.001909 |
| Left-vessel | 0.001129 |
| Left-Pallidum | 0.001000 |
| SupraTentorialVol | 0.000911 |
| BrainSegVol | 0.000812 |
| SupraTentorialVolNotVentVox | 0.000524 |
| SupraTentorialVolNotVent | 0.000481 |
| BrainSegVolNotVent | 0.000377 |
| BrainSegVolNotVentSurf | 0.000366 |
| Left-Hippocampus | 0.000329 |
| Right-Cerebellum-Cortex | -0.000447 |
| Left-choroid-plexus | -0.001078 |
| Left-Cerebellum-Cortex | -0.001522 |
| SubCortGrayVol | -0.001767 |
| SurfaceHoles | -0.001866 |
| CC_Posterior | -0.002304 |
| Right-Thalamus-Proper | -0.002689 |
| 5th-Ventricle | -0.003094 |
| 3rd-Ventricle | -0.004685 |
| Left-Thalamus-Proper | -0.005249 |
| Left-Inf-Lat-Vent | -0.005394 |
| Right-Putamen | -0.005477 |
| non-WM-hypointensities | -0.005570 |
| Right-Amygdala | -0.006690 |
| lhSurfaceHoles | -0.006709 |
| Left-Putamen | -0.007109 |
| BrainSegVol-to-eTIV | -0.007461 |
| Right-choroid-plexus | -0.007678 |
| Right-Hippocampus | -0.009197 |
| Optic-Chiasm | -0.010261 |
| Right-Pallidum | -0.010930 |
| Right-Cerebellum-White-Matter | -0.012078 |
| CC_Anterior | -0.013259 |
| lhCorticalWhiteMatterVol | -0.013764 |
| CorticalWhiteMatterVol | -0.013820 |
| rhCorticalWhiteMatterVol | -0.013856 |
| Left-Accumbens-area | -0.014673 |

## Notes

### Competing Interest Statement

The authors have declared no competing interest.

